# Novel kinome profiling technology reveals drug treatment is patient and 2D/3D model dependent in GBM

**DOI:** 10.1101/2022.07.22.499106

**Authors:** Federica Fabro, Nynke M. Kannegieter, Erik L. de Graaf, Karla Queiroz, Martine L.M. Lamfers, Anna Ressa, Sieger Leenstra

## Abstract

Glioblastoma is the deadliest brain cancer. One of the main reasons for poor outcome resides in therapy resistance, which adds additional challenges in finding an effective treatment. Small protein kinase inhibitors are molecules that have become widely studied for cancer treatments, including glioblastoma. However, none of these drugs have demonstrated a therapeutic activity or brought more benefit compared to the current standard procedure in clinical trials. Hence, understanding the reasons of the limited efficacy and drug resistance is valuable to develop more effective strategies toward the future. To gain novel insights into the method of action and drug resistance in glioblastoma, we established in parallel two patient-derived glioblastoma 2D and 3D organotypic multicellular spheroids models, and exposed them to a prolonged treatment of three weeks with temozolomide or either the two small protein kinase inhibitors enzastaurin and imatinib. We coupled the phenotypic evidence of cytotoxicity, proliferation, and migration to a novel kinase activity profiling platform (QuantaKinome™) that measured the activities of the intracellular network of kinases affected by the drug treatments. The results revealed a heterogeneous inter-patient phenotypic and molecular response to the different drugs. In general, small differences in kinase activation were observed, suggesting an intrinsic low influence of the drugs to the fundamental cellular processes like proliferation and migration. The pathway analysis indicated that many of the endogenously detected kinases were associated with the ErbB signaling pathway. We showed the intertumoral variability in drug responses, both in terms of efficacy and resistance, indicating the importance of pursuing a more personalized approach. In addition, we observed the influence derived from the application of 2D or 3D models in *in vitro* studies of kinases involved in the ErbB signaling pathway. We identified in one 3D sample a new resistance mechanism derived from imatinib treatment that results in a more invasive behavior. The present study applied a new approach to detect unique and specific drug effects associated with pathways in *in vitro* screening of compounds, to foster future drug development strategies for clinical research in glioblastoma.

## INTRODUCTION

Glioblastoma (GBM) is the most aggressive and frequent brain cancer in adults, with a median overall survival (OS) ranging between 14.6 and 16.7 months (1, 2). The standard treatment includes surgery, concomitant radiotherapy and chemotherapy with temozolomide (TMZ), followed by adjuvant administration of TMZ (1). One of the main problems of conventional therapy is that GBM becomes resistant in short term and, as a result after 6-9 months, the patient suffers from tumor recurrence (3, 4). This raises the urgency for a better understanding of the limited efficacy and resistance mechanisms occurring in GBM, in order to develop more effective strategies that overcome this problem.

At a molecular level, GBM is characterized by the presence of genetic alterations of molecules involved in crucial cellular functions such as proliferation, survival, invasion, altered metabolism, and evasion of immune response (5, 6). More specifically, alterations occur in protein kinases which are key regulators of canonical signal transduction pathways underlying these cellular functions. The most affected pathways in GBM are the RTK/PI3K/MAPK, p53, and Rb pathways, altered in 90%, 86% and 79% of cases, respectively (6). In the last decades, the aberrant phosphorylation activity of protein kinases has been a major target for anti-cancer treatments (7). As a result, the amount of small molecule protein kinase inhibitors (sPKIs) used as antineoplastic agents has significantly increased, with to date 55 out of 62 FDA approved sPKIs being used as antineoplastic agents (7, 8). For GBM treatment, several sPKIs targeting stem cells, growth, migration, cell cycle, cell death escape, and angiogenesis pathways have been tested for both newly diagnosed and recurrent GBMs (9-12). So far, however, sPKIs have shown limited efficacy in treating GBM as demonstrated by the failures of more than 100 clinical trials during the past twenty years (9-11). Among the promising small kinase inhibitors enzastaurin which is an inhibitor of the PKCβ and PI3K/AKT pathways, was tested up to a phase III trial for recurrent GBM, but failed to achieve superior efficacy compared to lomustine (13). Similarly, a limited antitumor activity was observed with imatinib. Imatinib is a multikinase inhibitor targeting PDGFR, ABL, c-KIT, that is already used for the treatment of gastrointestinal stromal tumor (GIST), and different types of leukemias (7). For GBM it had been tested for recurrent cases stopping at phase II clinical trial because of limited anti-tumor activity (14, 15).

One of the major reasons of these failures may be explained by the intrinsic or acquired tumor resistance developed to the compounds (10). A better understanding of the molecular mechanisms underlying the response to the drugs and their resistance has become a key point in improving GBM treatment. Nowadays, there is a growing need to increase the knowledge of tumor behavior and evolution under treatment (16). However, limited information is present regarding the pharmacodynamic effects of sPKIs that can explain their failure in GBM treatment. Known tumor-related resistance mechanisms of kinase inhibitors have been described for other cancers, potentially playing a role also in GBM. These include the acquisition of new mutations, co-activation of multiple oncogenic kinases, and activation of alternative signaling routes (17, 18). For brain tumors, additional lack of penetration and drug efflux pump activity in the blood brain barrier are also involved (19).

Preclinical *in vitro* screenings of compounds and drug candidates are largely used in drug discovery and development to understand their pharmacodynamic effects in tumor cells (20). An important factor that influences the therapeutic effect displayed *in vitro* is the dimensionality of the cell culture models used (21). Cell-based assays are still widely based on traditional two-dimensional (2D) cell cultures (20, 21). However, during the last 20 years, the use of three-dimensional (3D) cell models has exponentially increased (22). Recently, innovative 3D models in combination with organ-on-a-chip systems have been developed, introducing a more advanced representation of the tissue’s environment (23, 24). In addition, the latest development of mass spectrometry-based phosphoproteomics technology, which integrates high sensitivity and precision in the measurement of molecular targets, has resulted in its increased application to study mechanisms of action (MOA) and discover biomarkers directly, allowing a clearer understanding of mechanisms of resistance related to drug effects (25).

In this article, the molecular mechanisms of three drug candidates were investigated to improve the understanding of drug response in GBM. Two patient-derived GBM 2D cell cultures were established and treated with either TMZ, enzastaurin, and imatinib, to investigate the *in vitro* pharmacodynamic effects in GBM, and the heterogeneous response of the tumors after prolonged exposure. In parallel, we established fresh organotypic multicellular spheroids (OMS) to observe the influence of the dimensionality, both in suspended and organ-on-a-chip systems, on the outcome of the drug treatment. To better elucidate the MOA of the applied drugs, we coupled phenotypic evidence with the results of QuantaKinome™ analysis, a novel kinase activity profiling platform. This approach detects kinase activation loop phosphorylation status in a targeted manner and provides a more accurate and direct way to address endogenous pathway activity compared to conventional methods (25). Taken together, this study provides insights on the potential mechanisms of resistance that underly the failure of sPKIs for GBM treatment. We identified the heterogeneous activity of ErbB signaling pathway as the major player in 2D cell cultures, highlighting the impact of patient variability on medicine research. Furthermore, we charted the main phenotypic and molecular differences between 2D and 3D cell culture models, indicating another level of drug response variability to *in vitro* drug screenings. Additionally, this study underscores the relevance of combining phenotypic 2D or 3D model-derived evidence with kinase activity profiling, to perform functional studies which can offer novel strategic approaches for drug development and clinical research.

## MATERIAL AND METHODS

### 2D and 3D cell cultures models

Fresh glioma tissue samples GS.1012 and GS.1025 (Supplementary Table 1) were obtained directly from the operating room of the Erasmus Medical Center, the Netherlands. The use of patient tissue for this study was approved by the local Medical Ethical Review Committee Erasmus MC, code MEC-2013-090. Both patients signed an informed consent form according to the guidelines of the Institutional Review Board. The samples were taken directly from the operating theatre and placed in cold Dulbecco’s modified Eagle’s medium (DMEM/F12;Gibco) supplemented with penicillin and streptomycin (1%; Sigma). Within 2 hours post resection, samples were minced with surgical blades in small chunks. For each sample, the pieces were divided in two groups, one for 2D cultures and one for the 3D cultures. The 2D cell cultures were established in Dulbecco’s modified Eagle’s medium (DMEM)–F12 with 1% penicillin/streptomycin, B27 (Invitrogen), human epidermal growth factor (EGF; 20 ng/mL), human basic fibroblast growth factor (FGF; 20 ng/mL) (both from Tebu-Bio), and heparin (5 mg/mL; Sigma-Aldrich) by seeding the pieces in a basement-matrix-extracts (BME, Cultrex) coated petri dish. 3D organotypic multicellular spheroids (OMS) were created by adding each piece separately in a 96 well plate coated with 0.75% agarose (Invitrogen) to prevent attachment. After a week on average, the OMS acquired a spheroid shape. 192 and 128 OMS were generated for samples GS.1012 and GS.1025, respectively. All the cell culture models were cultured in serum-free condition as mentioned above.

### IC_50_ determination

To calculate the IC_50_ for temozolomide, enzastaurin, and imatinib, cells were seeded onto a basement membrane extract (BME, Cultrex)-coated 96-well plate at a density of 2000 cells/well. The plates were incubated for 24h prior to drug treatment. After 24h, serial 2-fold drug dilutions were prepared in serum-free culture medium and added to each well in technical triplicates. Ten concentrations of temozolomide (2.9 - 1500 μM), enzastaurin (0.1 – 100 μM) and imatinib (0.1 – 100 μM) were tested. The plates were incubated for additional five days. Viability was measured with CellTiter Glo 2.0 (Promega), a luminescent ATP assay, according to manufacturer’s instructions. The luminescence was measured with the Tecan Infinite F Plex. Percentage viability was normalized based on untreated controls.

### Drug treatment

Low passage (p2) 2D cell cultures were seeded in a 12-well plate in triplicate per condition, at a seeding density of 5000 cells/well. One half of the OMS (Supplementary Table 2) were placed, one per well, in a 384-well plate coated with 0.75% agarose (3D-SUSP). The other half (Supplementary Table 2) were placed in an organ-on-a-chip plate (Organoplate® Graft, Mimetas BV) (3D-OOAC). In short, 2 μl of ECM gel composed of 4 mg/ml type 1 collagen (Cultrex) was loaded into the ECM channel and allowed to polymerize at 37°C. Each OMS was placed in the grafting chamber to allow attachment. The plates were placed on an interval rocker (Perfusion rocker, Mimetas BV) set at a 7-degree inclination and 8-min cycle time.

Each drug-specific IC_50_ concentration was used to treat the 2D and 3D OMS cell cultures for three weeks. Specifically, the first treatment was performed three day after seeding. The weekly treatment regimen was carried out by culturing the cells in serum-free medium with drugs for three days, followed by refreshed medium without the drug for four days. 24 and 16 OMS were used per condition in GS.1012 and GS.1025, respectively.

### LDH and lactate assays

After the treatment, the cell culture medium was collected. LDH activity was performed using the LDH activity assay kit (Sigma-Aldrich) following the manufacturer’s instructions. The absorbance was measured at 450nm with the Tecan Infinite F Plex at 37°C every five minutes, for 26 cycles. LDH activity of the treatments samples were compared to the untreated controls.

Lactate concentration was measured in the cell culture medium using the LactateGlo assay (Promega) according to manufacturer’s instructions. Luminescence was measured with the Tecan Infinite F Plex. Lactate concentrations of the treated samples were compared to the untreated controls.

The LDH activity and lactate concentrations were normalized based on cells amount, OMS size, and OMS size with invasion area, in the 2D, 3D-SUSP and 3D-OOAC cell cultures, respectively.

### Scratch migration assay

Cells at a density of 10^4^ cells/well were plated onto BME-(Cultrex) coated 12-well plates in triplicate, and incubated at 37°C and 5% CO_2_. At confluency, monolayers were scratched with a P200 tip. Scratched monolayers were washed twice with sterile phosphate buffered saline (PBS). BME was added to cover the cells and allowed to polymerize at 37°C. Serum-free medium was added after 1h and phase-contrast pictures of the scratches were taken daily for 4 days using a phase contrast microscope (Observer D1, Zeiss). Cell migration was analyzed using ImageJ v.1.53 to measure the size of the wound, by averaging three measurements of the scratch at each time point. Data were expressed as percentage of the scratch area compared to 0 h.

### Cell counting and doubling time

Cells were seeded in an BME- (Cultrex) coated 96 well plate at a density of 1000 cells/well, in triplicate per condition and incubated at 37°C and 5% CO_2_. Cell counting was performed using a hemocytometer every 24 h for 7 days.

### Immunofluorescent staining

After the three weeks treatment, the medium of 3D OMS was replaced with new medium containing Calcein-AM (Thermo Fisher Scientific, diluted 1:2000) and Hoechst 33342 (Thermo Fisher Scientific, diluted 1:2000). The plates were incubated for 1 h at 37°C and 5% CO_2_. Images were acquired using a confocal microscope Micro XLS-C High Content Imaging Systems (Molecular Devices, US) at 4x magnification.

### Image acquisition

Phase contrast images of 3D OMS were acquired after each treatment using a microscope ImageXpress Micro XLS High Content Imaging Systems (Molecular Devices, US) at 4x magnification. The size of OMS was manually measured using ImageJ v1.53. The area of invading cells was calculated with ImageJ v1.53, after removing the background noise and adjusting the thresholds according to the cells invasion borders.

### Sample collection

After the three weeks treatment, samples were washed with cold PBS supplemented with phosphatase inhibitor (PhosSTOP, Roche) and protease inhibitor (cOmplete mini EDTA-free, Roche) and enzymatically dissociated with Accutase (Innovative Cell Technologies) or Dispase (1U/mL, Stemcell Technologies) at 37°C for 2D and 3D samples, respectively. OMS belonging to the same condition (22 and 15 OMS for GS.1012 and GS.1025, respectively) were pooled together prior the harvesting. Samples were centrifuged at 150g and washed with cold PBS supplemented with inhibitors twice. Finally, the pellets were frozen in liquid nitrogen. The samples were stored at -80°C.

### Kinase activity analysis using QuantaKinome™

To quantify human kinase activity in fresh frozen material, samples were processed and analyzed by Pepscope B.V. (Wageningen, the Netherlands). The QuantaKinome™ platform was applied to measure T-loop phosphopeptides by using a targeted LC-MS approach (QuantaKinome™, Pepscope B.V.). Briefly, frozen 2D and 3D samples were lysed and sonicated. For each sample, 100 μg and 44.7 μg aliquots of protein solution were processed for the 2D and 3D samples, respectively. After reduction, alkylation and digestion, all samples were dried and stored at -20°C until phosphoenrichment. Phosphorylated peptides were enriched and desalted using an automated platform. Samples were dried and stored at -80°C until LC-MS analysis. Next, samples within one experiment were measured in randomized order using the QuantaKinome™ targeted LC-MS assay (QuantaKinome™ Library v1, Pepscope).

Raw QuantaKinome™ data were analyzed by the manufacturer. The Principal Component Analysis (PCA) was performed to identify the variation between all samples in the data set. Multiple T-test analysis was used to determine log2 fold changes (2D and 3D samples) and to identify significantly regulated kinase activities (2D samples). Benjamini-Hochberg false discovery rate (FDR) < 0.05 was used to control for multiple testing. Pathway enrichment analysis was performed using the WikiPathways database (v10-01-2022) as pathway source.

### Data analysis

Statistical analyses were done using GraphPad Prism v8.4.2. For the dose-response data, nonlinear regression analysis was used to determine IC_50_ values. Doubling time was calculated using the exponential growth equation. Unpaired t-test was used to compare each drug treatment condition with the untreated control (LDH activity, lactate concentration, doubling time, OMS invasion). Repeated measure two-way ANOVA followed by Dunnett’s multiple comparisons test was used to compare the scratch migration between the drug treatments and control at each time point. Statistical significance was set at p < 0.05. Repeated measure two-way ANOVA followed by Tukey’s multiple comparisons test was used to compare the size of OMS for each condition at each time point. Statistical significance was set at p < 0.01. OMS data is presented as the mean ± standard error of the mean (SEM). 2D data is presented as the mean ± standard deviation (SD).

Network analysis was done using the STRING webtool v11.5 (www.string-db.org) using the multiple protein names search tool. The network settings were set to identify interaction with a minimum confidence score of 0.4 (medium confidence). The network was exported to Cytoscape v3.9.1.

## RESULTS

### Cytotoxicity, proliferation, and migration in 2D cultures of patient-derived glioblastoma stem-like cells (GSCs)

To determine the phenotypic effects of temozolomide (TMZ), enzastaurin (ENZA), and imatinib (IMA), two patient-derived GBM 2D cell lines (GS.1012 and GS.1025) were treated with their individual IC_50_ concentration (Supplementary Figure 1) for three weeks at intermittent intervals. The levels of cytotoxicity at the end of the three weeks were measured through the LDH activity present in the cell culture medium as a result of cell membrane damage. As shown in Figure 1A, the treatments differentially affected the two GBMs.

**Figure 1.**
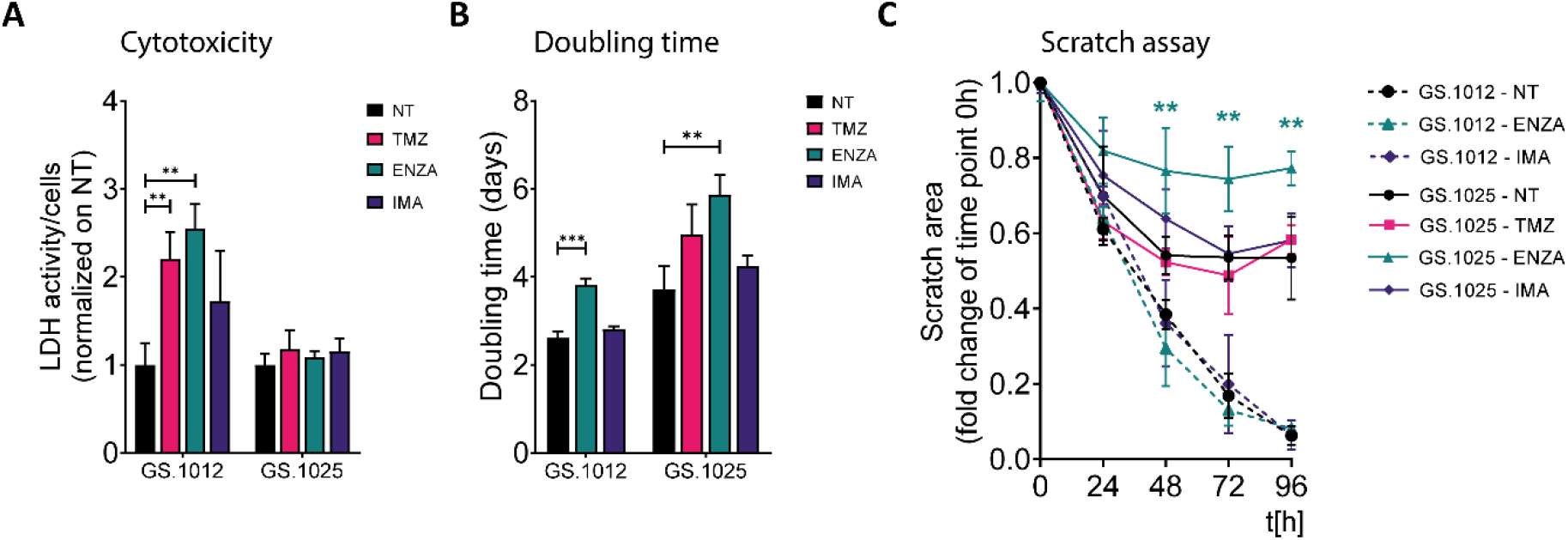
LDH activity, proliferation, and migration assay of GS.1012 and GS.1025 2D cell cultures. (A) Bar plots show the LDH activity in samples GS.1012 and GS.1025 for temozolomide (TMZ), enzastaurin (ENZA) and imatinib (IMA) treated samples normalized to values observed in untreated (NT) condition. (B) Bar plots show the cell proliferation expressed as doubling time in days, of GS.1012 and GS.1025 in untreated (NT), temozolomide (TMZ), enzastaurin (ENZA) and imatinib (IMA) conditions. (C) Line plots show the percentage of the scratched area (normalized to values observed at t=0h) across time (hours). Continuous lines represent GS.1012, while segmented line represents GS.1025. Data are presented as mean ± SD. *p < 0.05; **p < 0.01; ***p<0.001.

Particularly, an increased cytotoxicity compared to the untreated condition was detected in the GS.1012, where toxicity levels were more than doubled after exposure to TMZ and ENZA (2.2- and 2.5-fold increase, respectively). In contrast, the levels of LDH activity in GS.1025 remained stable for all treatments. To extend the investigation on the toxicity, the concentration of lactate in the supernatant was measured as a marker of cell stress in *in vitro* repeat-dose testing regimes (26). The concentrations of lactate confirmed the elevated cellular stress derived from the TMZ and ENZA treatments in GS.1012, and the lack of toxicity in GS.1025 (Supplementary Figure 2). As a consequence of the heavy toxicity throughout the treatment, the TMZ treated cells of GS.1012 could not be recovered and further tested, confirming its potency in GBM.

Given that proliferation and migration are relevant cellular functions in cancer progression, and important targets to assess treatment efficacy (27), the impact of TMZ and sPKIs treatment on these two behaviors was assessed. The response to proliferation was derived by calculating the cell doubling time. As shown in Figure 1B, ENZA was the only compound that impacted the proliferation of cells in both patients, displaying a significant 1.4- and 1.6-fold increase of the doubling time in GS.1012 and GS.1025, respectively. Consistent with the inefficacy observed in the cytotoxicity assays, imatinib did not affect the proliferation of the cells. In GS.1025, TMZ showed a trend in slowing down the proliferation. To test the migration ability, a scratch migration assay was performed (Figure 1C, Supplementary Figure 3). Overall, none of the treatments had impact on the migrative trait of GS.1012. Interestingly, GS.1025 was characterized by a low migratory ability. Only 53% of the gap was closed in the untreated condition. Nevertheless, ENZA further affected the poor migration ability of these cells. After 96 hours, the migration was significantly reduced by 45% compared to the untreated condition.

Overall, these results showed that TMZ was the only compound that could eliminate the entire tumor population in at least one of the patients samples, demonstrating its higher potency compared to the two sPKI. Regardless of the promising effects of ENZA treatment in reducing proliferation and migration, its overall effect did not eradicate completely the tumor cells in both patients, indicating the presence of counteracting resistance mechanisms. Interestingly, in contrast to ENZA and TMZ, IMA had no cytotoxic effects suggesting a strong intrinsic resistance of both GBMs to the compound.

### Protein kinase activity in 2D cell cultures of patient-derived GSCs

To investigate the molecular MOA and potential resistance response of TMZ, IMA, and ENZA treatment, the QuantaKinome™ platform has been used. This technology is capable of measuring the intracellular protein kinase activity by quantifying the activation-loop phosphorylation of endogenous kinases and provides a functional profile of the sample (25). A total of 55 and 63 different kinase activities were quantified in GS.1012 and GS.1025, respectively, covering all major kinases groups (Figure 2A, Supplementary Figure 4).

**Figure 2.**
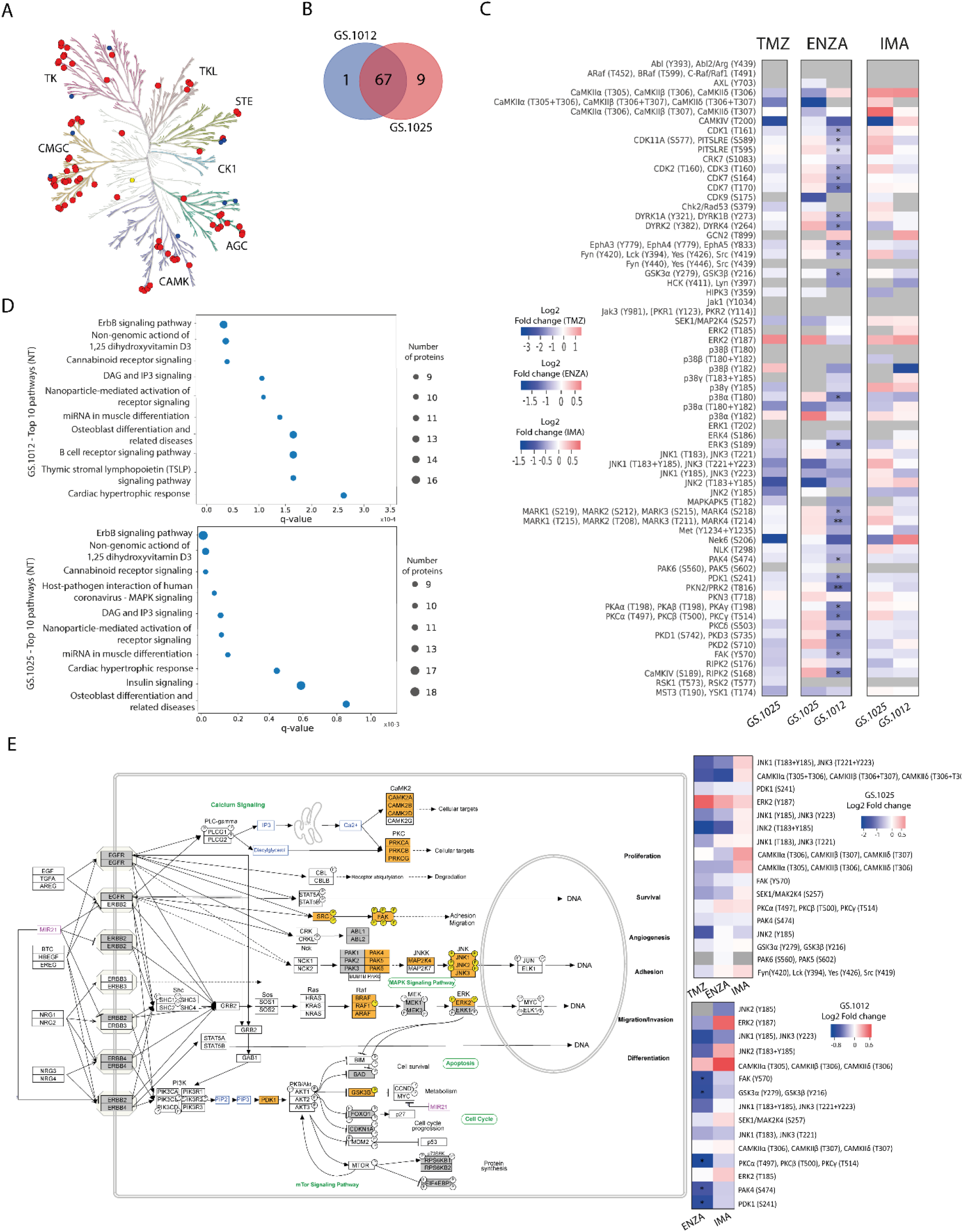
Kinase activity and pathway analysis of GS.1012 and GS.1025 2D cell cultures. (A) Human kinome tree representing the kinase groups detected with QuantaKinome™. Red dots represents the kinases detected in both GS.1012 and GS.1025 samples. Blue dots represent kinases detected only in GS.1025. Yellow dots represent kinases detected only in GS.1012. (ACG: Containing PKA, PKG, PKC families; CAMK: Calcium/calmodulin-dependent protein kinase; CK1: Casein kinase 1; CMGC: Containing CDK, MAPK, GSK3, CLK families; STE: Homologs of yeast Sterile 7, Sterile 11, Sterile 20 kinases; TK: Tyrosine kinase; TKL: Tyrosine kinase-like). Illustration made with KinMap (www.kinhub.org/kinmap/). (B) Venn Diagram showing the number of common and specific kinases identified in GS.1012 and GS.1025. (C) Differential activity analysis in 2D cell cultures treated with temozolomide (TMZ), enzastaurin (ENZA), and imatinib (IMA) in GS.1012 and GS.1025. Heatmaps showing the log2 fold changes of kinase activity of the treatments compared to the untreated controls. Gray boxes represent kinase activities identified in the sample for either the treatment or the control. *FDR adjusted-p <0.05; ** FDR adjusted-p <0.01. (D) Pathway analysis of 2D GS.1012 and GS.1025 cell cultures. Dotplot showing the top 10 pathways using the detected kinases in GS.1012 (top) and GS.1025 (bottom) untreated cells. The dot size represents the number of kinases detected, while the distribution on the x-axis indicate the q-value. The pathways source is WikiPathways. (E) ErbB signaling pathway representation according to WikiPathways of kinases detected in GS.1012 and GS.1025. The orange boxes represent the detected kinases with QuantaKinome™. The pathway has been acquired from WikiPathways (www.wikipathways.org/instance/WP673). The gray boxes represent kinases present in the QuantaKinome™ library but not detected in the study. On the right, the heatmaps showing the log2 fold changes of the kinases belonging to the Erbb signaling pathway in GS.1025 (top) and GS.1012 (bottom).

Of these kinase activities, 87% (n=67) was detected in both GBM samples, while 1.3% (n=1) was specific of GS.1012, and 11.7% (n=9) was specific of GS.1025 (Figure 2B). The detected kinases comprised relevant members involved in the aberrant signaling networks of GBM, such as MAPKs, Src family, PKCs, CDKs, CAMK2s, and are here used to identify treatment specific response profiles to capture the MOA of the drugs.

For each patient separately, an unsupervised hierarchical clustering analysis was performed using all the quantified kinase activities to test the similarity of treatments response. As expected, the different treatments grouped separately from each other indicating that kinase activity response was treatment specific (Supplementary Figure 5). The TMZ and ENZA treatment replicates clustered together suggesting a homogeneous response. IMA replicates, however, clustered more closely to each other or with the untreated samples. Interestingly, the untreated conditions displayed a more heterogeneous molecular profile, reflecting the intratumoral heterogeneous nature of treatment naive GBM.

To identify the kinases most affected by the treatments, and to achieve a better understanding of the MOA, a differential kinase activity analysis was performed. In contrast to TMZ toxicity that caused the death of GS.1012 cells, in GS.1025 the treatment did not induce significant changes in the activation of the measured kinases (Figure 2C). Nevertheless, TMZ induced a shift towards the inhibition of the intracellular signaling cascade regulated by JNK kinases, up to CAMK4 and NEK6, which are regulators of transcription and cell cycle, respectively (28, 29). In contrast, the mitogenic ERK2 displayed an increased activity, indicating a sustained activation of a survival signal.

The specific target of the sPKI ENZA is PKCβ. As shown in Figure 2C, a significant downregulation of PKCs activation was measured in GS.1012 treated with ENZA. Interestingly, the inhibition of PDK1, an upstream regulator of PKC, was also observed. As a result, we identified in GS.1012 a significant downregulation of the direct downstream molecules of both PDK1 and PKC such as PKA isoforms, GSK3α, GSK3β, FAK, PKD1, PKD3, and PKN2 (Figure 2C). In addition, other kinases activated by PKC include also the Src family comprising LCK, LYN, SRC, and YES. These kinases were also affected reaching, as final targets, kinases strictly linked to cellular proliferation and migration. This was demonstrated by the significant inhibition of the cell cycle cyclin-dependent kinases (CDKs) like CDK1, CDK2, CDK3, CDK7, and CDK11, and the decreased FAK and ERK3 activity, which both play a role in cell migration in cancer (30-33). To confirm the relation between ENZA targets and the significantly affected kinases, an interaction network analysis was performed with STRING for GS.1012 sample. As shown in Supplementary Figure 6A, there is a highly interconnected association of ENZA targets with the significantly impaired downstream kinases, displaying the overall affected intracellular network. In GS.1025, PKCs activity was not affected by ENZA, while PDK1 showed only a modest trace (log2FC=-0.21) of decreased activity (Figure 2C). Moreover, the downstream ERK2 and the cell cycle CDKs displayed a sustained activation of proliferation signals. Nevertheless, an inhibitory tendency affecting the MAPKs MAP2K4, p38α and JNKs was seen, which could have led to the observed proliferative slowdown. In addition, the results showed a trending dysregulation of CDK9, involved in the transcriptional regulation, and MST3 and YSK1, which are kinases involved in the regulation of cell polarity, adhesion, and migration (34-36). Due to their role, these two kinases could have furtherly influenced the disruption of the migration ability observed in the scratch migration assay.

As a multikinase inhibitor, IMA was expected to target PDGFR, KIT, and ABL. However, none of the IMA targets were endogenously detected in the samples. Nevertheless, in accordance with the absent cytotoxicity, and sustained proliferation and migration, we did not observe significant changes in kinases activity (Figure 2C). In both tumors, a sustained or increased activation trend was observed for relevant kinases involved in survival and proliferation such as the members of the MAPK kinases family MAP2K4, ERK2, p38γ, and JNK2. In GS.1025, additional proliferative support signals were derived from the intermediaries FYN, SRC, LCK, YES, and the cell cycle regulating kinases CDK1, CDK2, CDK3, CDK7, CDK11, and CHK2. Moreover, we identified other kinases with higher but not significant activation such as PKAs, PKCs, MARKs, DYRK1A, DYRK1B, CDK9, and PKN3. The network analysis revealed that IMA targets are strongly associated with the quantified downstream kinases that display a sustained activity after the treatment (Supplementary Figure 6B), indicating the inefficacy of IMA in affecting the downstream signaling processes.

For a clearer understanding of the intracellular signaling events after the treatments, a pathway enrichment analysis was performed using the detected kinases for both patients. The results indicated ErbB signaling as the most representative pathway across conditions (Figure 2D, E). As shown in Figure 2E, ErbB receptors activate a multiplicity of downstream intracellular signals that regulate crucial cellular processes in cancer such as proliferation, survival, migration, invasion, angiogenesis, adhesion, and differentiation (37). Of the identified ErbB signaling pathway kinase activities in GS.1012, 33.3% (5/15) were significantly reduced by ENZA, 33.3% (5/15) displayed a trending inhibition, and the remaining kinases showed no changes or a trending increase (Figure 2E). On the other hand, in GS.1025, 41.1% (7/17) showed a trending inhibition, while the remaining 58.9% (10/17) showed no changes or a trending increase. With IMA treatment, 53.3% (8/15) of the ErbB kinases in GS.1012 displayed a trending decreased activity, while the remaining 46.7% (7/15) displayed no changes or a trending increase (Figure 2E). In GS.1025, only 23.5% (4/17) of the kinase activity showed a trending decreased, while the remaining 76.5% (13/17) showed no changes or a trending increase.

Overall, the findings derived from the drug treatment analyses conducted on 2D patient-derived GSCs revealed the inter-tumoral variability of the intracellular signals in response to drug treatments. In the specific case of TMZ, despite displaying a trending inhibition, the prolonged treatment did not induce a sufficient suppression of kinases activity in the surviving tumor cells. The results from the sPKIs treatment indicated that ENZA caused the increased cytotoxic levels when it successfully inhibited its targets, resulting in a significant inhibition of CDKs through the MAPKs cascades. On the other hand, IMA had less impact due to the constant MAPKs activation that played a central role in sustaining the survival signals in both tumors.

### Cytotoxicity, proliferation, and migration in 3D OMS of patient-derived GSCs

To investigate the differences in drug response between 2D and 3D cell culture models, 3D organotypic multicellular spheroids (OMS) derived from the patients GS.1012 and GS.1025 were generated in parallel. The spheroids were seeded in a low attachment environment (3D-SUSP), and in an organ-on-a-chip platform (3D-OOAC) containing a matrix of collagen 1 to additionally examine the effects of 3D OMS on migration. The OMS were treated with TMZ, ENZA, and IMA with the same treatment regimen and concentrations used for the 2D cell cultures.

To understand the degree of cytotoxicity developed in the two 3D models, the LDH activity and lactate levels were measured. In GS.1012, in contrast to the 2D cultures, the OMS were not affected by the treatments. In fact, the LDH activity of the 3D-SUSP was significantly reduced compared to the untreated condition by 3.1-, 4.0-, and 14.3-fold decrease in TMZ, ENZA, and IMA treatments, respectively (Figure 3A).

**Figure 3.**
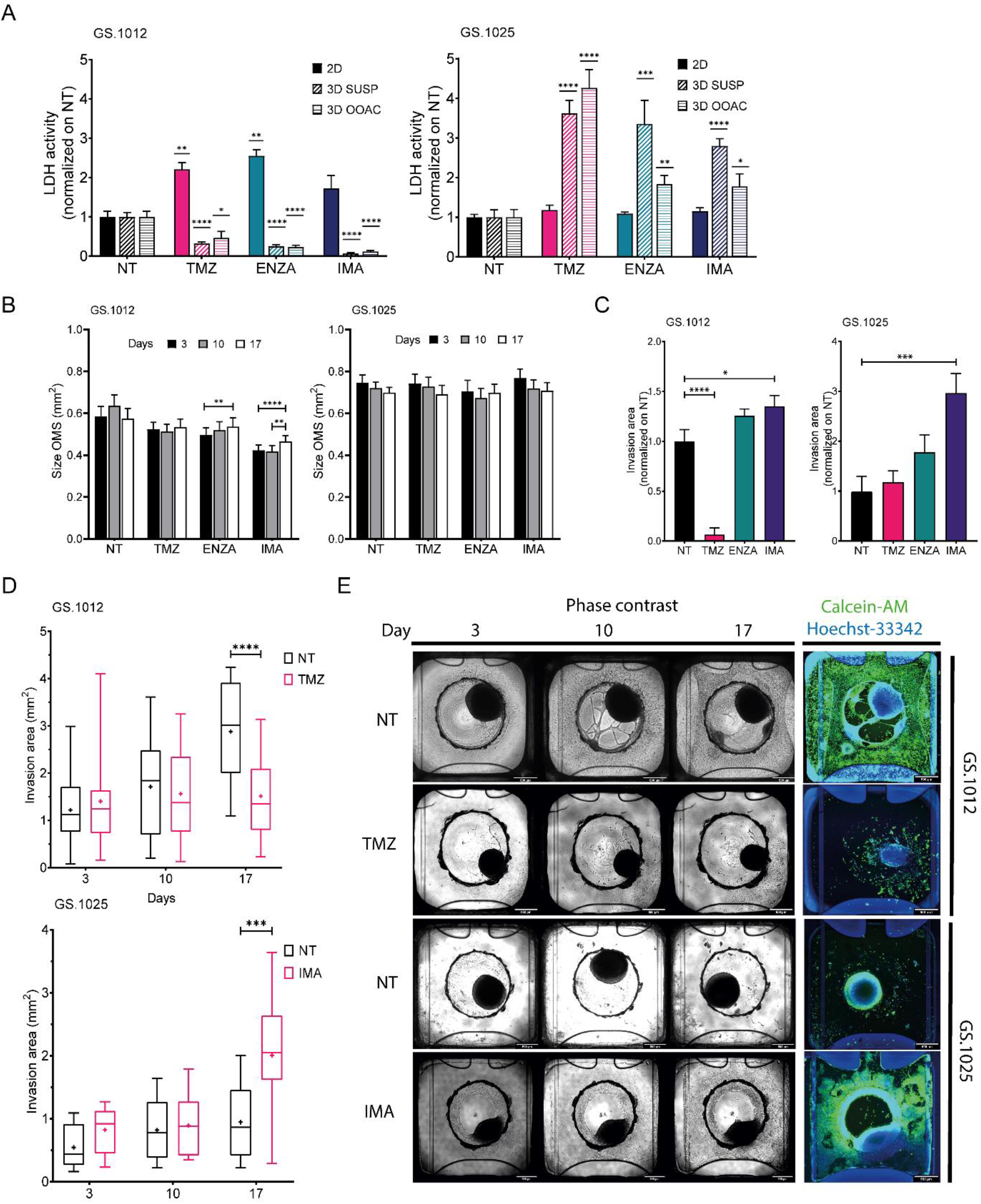
LDH activity, size, and migration of GS.1012 and GS.1025 3D cell cultures. (A) Bar plots showing the LDH activity in 2D, 3D-SUSP and 3D-OOAC samples of GS.1012 (left panel) and GS.1025 (right panel) in temozolomide (TMZ), enzastaurin (ENZA) and imatinib (IMA) treated samples normalized to values observed in non-treated (NT) control. n(GS.1012)=22; n(GS.1025)=16. Data are presented as mean ± SD. *p < 0.05; **p < 0.01; ***p <0.001; ****p <0.0001. (B) Bar plots showing the size variation in time of 3D-SUSP OMS of GS.1012 (A) and GS.1025 (B) samples in non-treated (NT), temozolomide (TMZ), enzastaurin (ENZA) and imatinib (IMA) treated conditions. n(GS.1012)=22; n(GS.1025)=16. Data are presented as mean ± SEM. **p < 0.01; ***p <0.001; ****p<0.0001. (C) Bar plots showing the invasion area of 3D-OOAC OMS of GS.1012 (A) and GS.1025 (B) samples in temozolomide (TMZ), enzastaurin (ENZA) and imatinib (IMA) treated conditions normalized on non-treated (NT) control. n(GS.1012)=22; n(GS.1025)=16. Data are presented as mean ± SEM. *p<0.05; ***p <0.001; ****p<0.0001. (D) Invasion area of 3D-OOAC OMS through treatment time in GS.1012 and GS.1025. Box plots show the distribution of the invasion area of 3D-OOAC OMS of GS.1012 (top panel) and GS.1025 (bottom panel) through treatment time, in untreated (NT), temozolomide (TMZ), enzastaurin (ENZA) and imatinib (IMA) conditions. The middle line represents the median. The cross represents the mean. n(GS.1012)=22; n(GS.1025)=16. ***p <0.001; ****p<0.0001. (E) Development of OMS in the Organoplate®Graft. Representative images show GS.1012 and GS.1025 OMS in the Organoplate®Graft after 3, 10, and 17 days in untreated (NT, top) and temozolomide (TMZ, bottom) conditions. On the right, the OMS are stained with Calcein-AM to identified the living cells.

A similar trend was observed in the 3D-OOAC, where the toxicity was 2.1, 4.3, and 8.6 times lower than the untreated condition (Figure 3A). Compared to corresponding 2D cell cultures, the OMS displayed more resistance to the therapies, with IMA as the least effective drug as indicated by the lowest LDH activity. The lactate concentrations confirmed the absence of therapy-related toxicity except in the 3D-OOAC samples treated with TMZ, which displayed the highest LDH activity among the treatments (Supplementary Figure 7A). Interestingly, opposite to GS.1012, the 3D OMS of GS.1025 displayed more sensitivity compared to their 2D counterpart. The LDH activity measured in the 3D-SUSP GS.1025 samples was 3.6-, 3.3-, and 2.8-fold higher than the untreated condition in TMZ, ENZA, and IMA treatments, respectively (Figure 3A). Similarly, the 3D-OOAC displayed a significant LDH activity increase of 4.2, 1.8, and 1.7-fold of the untreated conditions. In contrast to the LDH activity, the lactate concentration did not indicate the presence of toxicity, except for 3D-SUSP treated with ENZA (Supplementary Figure 7B).

To assess the drug effects on proliferation and migration in 3D setting, we evaluated the changes in spheroids size and matrix invasion in 3D-SUSP and 3D-OOAC, respectively. Significant but not relevant increases in size were seen in GS.1012 3D-SUSP treated with ENZA and IMA, while no significant differences were observed in GS.1025 (Figure 3B). In the 3D-OOAC, the migration ability was drastically reduced in GS.1012 treated with TMZ by 15.8 times, while it was slightly increased after IMA treatment (Figure 3C-E). Consistent with its 2D counterpart, GS.1025 3D OMS display a low level of invasiveness under control conditions (Figure 3D). Interestingly, IMA significantly induced an invasive phenotype by increasing the invasion ability by 3-fold (Figure 3C-E).

Overall, these results indicated that in both the 3D OMS types of both patients, despite the opposite cytotoxic measurements, TMZ displayed the highest levels of cytotoxicity compared to the other drugs. TMZ appeared to be a potent drug, particularly in monolayer or exposed invading cells, while the 3D architecture appears to confer protection to the cells. Opposite to TMZ, IMA was found to be the least effective compound, even inducing a more invasive phenotype. A general lack of efficacy to IMA was observed also in 2D cell cultures, which however did not display more aggressive traits as the enhanced migration ability. Consistently, ENZA demonstrated to be more effective in 2D cell cultures, while 3D OMS were not significantly affected.

### Protein kinase activity in 3D OMS cell cultures of patient-derived GSCs

To gain an insight on the influence of the 3D architecture on treatment response and kinase activation, the OMS of each condition were collected to perform QuantaKinome™ analyses. A total of 53 and 59 different kinase activities were quantified in GS.1012 and GS.1025, respectively (Supplementary Figure 8). The principal component analysis (PCA) and the hierarchical clustering heatmaps showed a separation in kinase activity between 2D and 3D cell cultures along PC1 (Figure 4A, Supplementary Figure 9).

**Figure 4.**
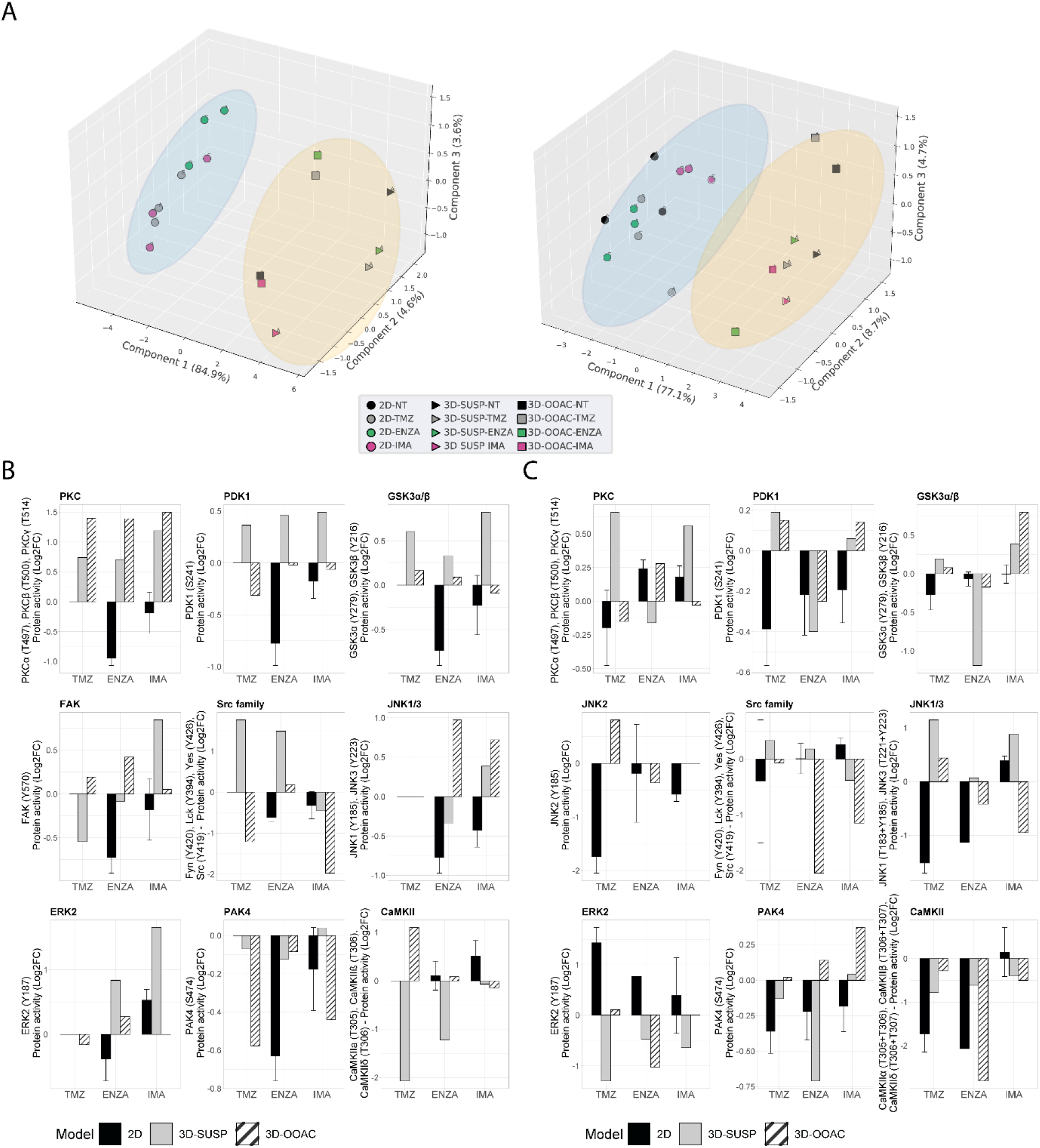
Comparison of kinase activity of GS.1012 and GS.1025 2D and 3D OMS cell cultures. (A) Principal component analysis (PCA) of kinase activities. 3D principal component analysis performed on the commonly identified kinase activities in 2D, 3D-SUSP, and 3D-OOAC of untreated (NT), temozolomide (TMZ), enzastaurin (ENZA), and imatinib (IMA) conditions of GS.1012 (left) and GS.1025 (right). The blue circle clusters the 2D samples, while the yellow circle clusters the 3D OMS samples. (B) Kinase activity of Erbb signaling kinases in 2D and 3D OMS of GS.1012. The barplots show the log2 fold changes values of detected kinases belonging to the Erbb signaling pathways that were identified in 2D, 3D-SUSP, and 3D-OOAC of temozolomide (TMZ), enzastaurin (ENZA), and imatinib (IMA) conditions normalized to the untreated (NT) control. n(2D)=3 (except JNK1/3-ENZA and CAMK2-IMA: n=2); n(3D)=1;* FDR adjusted-p < 0.05; ** FDR adjusted-p <0.01. (C) Kinase activity of Erbb signaling kinases in 2D and 3D OMS of GS.1025. The barplots show the log2 fold changes values of detected kinases belonging to the Erbb signaling pathways that were identified in 2D, 3D-SUSP, and 3D-OOAC of temozolomide (TMZ), enzastaurin (ENZA), and imatinib (IMA) conditions normalized to the untreated (NT) control. n(2D)=3 (except ERK2-IMA, JNK1/3-TMZ, JNK2-ENZA, JNK2-IMA: n=2; ERK2-ENZA, JNK1/3-ENZA, CAMK2-ENZA: n=1) n(3D)=1; *FDR adjusted-p < 0.05; ** FDR adjusted-p <0.01.

A specific group of kinases displayed an higher activity in both GS.1012 and GS.1025 3D cultures compared to the 2D: CAMK2s, ERK3, and ERK4 (Supplementary Figure 9). To note, the identified CAMK2 activities at T305/306 are inhibitory (38). Furthermore, the 3D-OOAC appeared to be closer to the 2D cultures compared to the 3D-SUSP in GS.1012, indicating the presence of similar features that could derive from the migrative ability. In addition, a further separation along PC1 could be observed between the distributions of 3D-SUSP and 3D-OOAC in GS.1012, while in GS.1025 they appeared more homogeneous.

Next, to explore the influence of the 3D models on the ErbB signaling pathway, an initial observational analysis of the main differences of the related kinase activity between 2D and 3D OMS was carried out. As shown in Figures 4B-C, we observed a diverse spectrum of kinase regulation patterns after the different treatments.

With regard to the significantly reduced invasion in TMZ treated 3D-OOAC of GS.1012, the contribution of the ErbB signaling pathway is suggested to derive from the enhanced CAMK2 inhibition (log2FC: 1.10), and from a decreased Src family (log2FC: -1.21), PDK1 (log2FC: - 0.31), ERK2 (log2FC: -0.16) activity (Figure 4B). The kinases involved in the ErbB signaling mostly displayed an increased activity in 3D-SUSP OMS, with the exceptions of Src family and FAK kinases (Src family log2FC: -1.21; FAK log2FC: -0.54) (Figure 4B). These kinases are involved in the adhesion and migration, and their reduced activity reflects the missing migrative feature of the 3D-SUSP OMS.

In both 3D cultures of GS.1012, consistent with the reduced LDH activity, the main targets PDK1 and PKCs were not affected by ENZA. In contrast to what resulted in the 2D cell cultures, their activity was constant (PDK1 3D-OOAC log2FC: -0.02) or increased (PDK1 3D-SUSP log2FC: 0.46; PKCs 3D-SUSP log2FC: 0.70; PKCs 3D-OOAC log2FC: 1.39) compared to untreated condition (Figure 4B). A tendency of upregulating the downstream kinases was also observed, derived in particular from GSK3α, GSK3β, Src family, and ERK2 in both 3D cell culture models (Figure 4B).

IMA, which displayed the least cellular toxicity in both 2D and 3D cell cultures, showed a constant or enhanced activity of most kinases of the 3D OMS compare to the untreated condition, in contrast with the 2D cultures. In the 3D-SUSP, the downregulation of kinase activity was restricted mainly to Src family (log2FC: -0.45), while in the 3D-OOAC it was displayed more intensely by both the Src family kinases and PAK4 (Scr family 3D-OOAC log3FC: -1.97; PAK4 3D-OOAC log2FC: -0.44). Interestingly, the activity of these kinases was reduced also in the 2D cultures (Figure 4B). Overall, consistently to the decreased levels of cellular toxicity observed in GS.1012, we found mainly a sustained or increased activation of crucial players in the control of growth and proliferation, linked to PDK1, PKCs, GSK3A, GSK3B, FAK, JNK1/3, and ERK2.

In GS.1025, an overall diminished mitogenic ERK2 activity was observed in the 3D OMS treatments (Figure 4C), in line with the increased cytotoxicity compared to the 2D cultures. After TMZ treatment, however, an increased activity of JNKs (JNK1/3 3D-SUSP log2FC: 1.15; JNK1/3 3D-OOAC log2FC: 0.44; JNK2 3D-OOAC log2FC: 0.81) was measured, indicating an alternative survival signal than ERK. In fact, JNKs, defined as stress-activated MAPKs, orchestrate cellular responses to many types of stresses and promote survival in cancer (39). Moreover, it has been associated with enhanced TMZ resistance in GBM (40).

Interestingly, ENZA appeared to reduce to a higher extent the kinases involved in the ErbB signaling in the 3D OMS than in 2D cell culture, as an indication of the higher levels of cellular toxicity observed compared to the 2D cell cultures. In particular, PDK1 was affected by the treatment in both 3D OMS cultures (3D-SUSP log2FC: -0.40; 3D-OOAC log2FC: -0.39), while PKCs activity was slightly reduced only in the 3D-SUSP OMS (3D-SUSP log2FC: - 0.16). In addition, a downregulation of the kinase activity was observed particularly for GSK3A, GSK3B, PAK4, Src family, ERK2, and CAMK2 in either one or both 3D culture models.

Finally, of notice, IMA treatment induced the activation of GSK3α and GSK3β (3D-SUSP log2FC: 0.39; 3D-OOAC log2FC: 0.40), which was observed only in 3D cultures of the IMA-treated samples, and, with a higher extent, compared to the other drugs (Figure 4C). A common tendency behavior in both the 3D OMS cultures that differed from the 2D counterpart was observed with the increased activity of PDK1 and PAK4, and a reduced activity of the Src family kinases.

## DISCUSSION

Limited drug efficacy and resistance are major issues in treating GBM. While for several cancers small molecule protein kinase inhibitors have successfully brought a therapeutic benefit, this has not been achieved for GBM as demonstrated by the amount of unsuccessful clinical trials (11). Reaching a better understanding of the mechanisms of action (MOA) taking place during drug treatment can shed light on the reasons why these promising compounds alone did not succeed, and guide future drug development in a more successful direction.

Our study is the first to investigate the drug response in patient-derived cell cultures by coupling the phenotypical evidence with QuantaKinome™, a novel molecular functional platform that measures the activity of endogenous kinases crucial in cancer. Our approach allows the identification of key signaling pathways that contribute to the efficacy and toxicity, as well as to the resistance, occurring after drug treatments. In fact, nowadays it has become of crucial importance to understand the MOA of the drugs at a molecular level, and identify potential pitfalls of novel therapies before reaching the clinical phases. Our results highlight three major aspects contributing to the tumor resistance to sPKIs and consequent failure of these drugs in the treatment of GBM: the inter-tumoral heterogeneity, the presence of constantly active bypass signals derived mainly from the activation of the ErbB pathway, and the influence of the preclinical evaluation of the drugs in different model systems. The first two aspects were investigated primarily on the 2D cell cultures. Due to the heterogeneous nature of GBM, we observed different phenotypical behaviors and kinase activities between patients, particularly under TMZ and ENZA treatments.

TMZ, as an alkylating agent, does not target specific kinases. The alterations of the intracellular signaling are secondary effects derived from the DNA damages. The survival signaling that was measured in the surviving TMZ treated cells was derived in particular from a trending upregulation of ERK2. In fact, MAPKs are considered key transducers of aberrant signaling in GBM and their downregulation has proven to be a marker of induced TMZ toxicity (40-43).

The sPKI ENZA mostly impaired the viability of tumor cells when it significantly inhibited its main target PKCβ. Nevertheless, in parallel, we also observed the downregulation of the other PKC isoforms and PDK1, an upstream regulator of PKC, followed by the downregulation of the signaling cascade, through Src family kinases up to CDKs. In a previous study it was demonstrated that ENZA has a modest activity also on PDK1 which could explain the inhibition measured (44). Moreover, PDK1 represents a divergence point for receptor tyrosine kinases, such as ErbBs, and the regulation of multiple signaling cascades including the activation of PKC (44-46). Another significant downregulation of kinase activity downstream of PDK1 was observed for GSK3α and GSK3β kinases. In line with our observation, the downregulation of GSK3β activity is considered a reliable pharmacodynamic marker for ENZA efficacy (47). ENZA resistance has not been studied in GBM. In our study, the resistance to ENZA was more evident in one of the two cell cultures, and was characterized by the unsuccessful inhibition of PKC, PDK1, GSK3α and GSK3β activities, and a concomitant increased trend of p38α and ERK2. A phase III clinical trial of ENZA combined with TMZ and radiation therapy in patients with newly diagnosed GBM is currently ongoing (NCT03776071).

In contrast to TMZ and ENZA, IMA showed a general lack of efficacy in both GBM samples. However, the kinomic profile observed after the treatment with IMA was different between the two GBMs, indicating the heterogeneity of intrinsic resistance mechanisms. Imatinib resistance has been mainly studied for chronic myeloid leukemia (CML) and GIST, where the most common cause is the development of new mutations in the kinase domains (48-50). Nevertheless, a study identified also a persistent activation of Fyn/ERK signaling in imatinib-resistant leukemia cells (51). Similarly, we detected the sustained activation of Fyn/ERK2 in GS.1025 and ERK2 in GS.1012, also suggesting that resistance is not dependent only on the effects on the main targets.

Many of the kinases identified in our study are important members of the ErbB signaling pathway, which is responsible for the stimulation of several key interconnected intracellular signals and proteins that promote tumor survival and progression (37). In 2D cell cultures, we observed a heterogeneous response of ErbB pathway kinases to drug treatments in terms of activity. Clear evidence is seen with ENZA treatment, where GS.1012 showed a significant inhibition of ErbB pathway kinases that was not observed in GS.1025. Another case is represented by IMA treatment, where GS.1025 displayed a trending increased activity of most kinases, in contrast with GS.1012. In addition, we identified ERK2, of which activity showed a trending enhancement common to all the drug treatments of GS.1025. While in TMZ treatment ERK2 appeared to play a more relevant sustaining role of the pathway, in ENZA and IMA treatments a trending activation was measured also for other members of the pathway. Due to its mitogenic function, ERK activity indicates a sustained activation of survival signals and, therefore, provides an open door to the sustainment of resistance. As key players in the ErbB signaling pathway, ERK kinases have been correlated with tumor progression, shorter overall survival, and higher proliferation indices in GBM (52, 53). As such, ERK2 activity represent a potential target to pursue for GBM treatment and to overcome resistance in a personalized context. ERK inhibitors have already been developed for other cancers and are being currently tested in clinical trials in combination with other drugs in GBM, representing a promising target for GBM therapy (11, 54).

The second part of our study focused on the influence derived from different cell culture models in the investigation of the sPKIs’ MOA and resistance to these treatments. It is beyond the scope of this study to state which model is the best to assess the efficacy of the drugs, as the selection of the most relevant model depends on the aim of the study. Drug screenings in 2D and 3D GBM cell cultures have already been investigated (55). However, our approach provides additional relevant information through the application of organ-on-a-chip system, which is a technology that can be exploited to recreate multiple environmental interactions, and the direct read-out of kinase activity (25, 56). Due to the lack of replicates and low amounts of proteins for 3D OMS, we limited our analysis on ErbB signaling kinases that were identified in common between the 2D cell cultures. Nevertheless, we could observe some relevant phenotypical and molecular differences in response to the drugs between 2D and 3D cell cultures, and, to a lesser extent, between 3D-SUSP and 3D-OOAC, derived from the different invasive potential. The concomitant treatment of these three different cell culture models provided additional information on the drugs responses. This was particularly evident with the GS.1012 TMZ-treated and GS.1025 IMA-treated samples. In fact, TMZ displayed its maximal and minimal effect on the 2D and 3D-SUSP cultures, respectively, and partial efficacy in the 3D-OOAC derived from the impaired migration. All the results together indicated that only directly-exposed cells were sensitive to the toxic effects of TMZ, suggesting a protective role derived from stroma and the cell-cell interactions of the 3D structure. Indications of toxicity at kinomic level in the 3D-OOAC were linked to the enhanced inhibition of CAMK2 inhibitory activity, as a potential consequence of the decreased calcium signaling, and a reduced activity of PDK1, Src family, and ERK2 kinases, as a signal of cellular stress.

The inefficacy of IMA in GS.1025 was observed both in 2D and 3D OMS. Nevertheless, in the 3D-OOAC cultures IMA greatly induced migration, proving not only the resistance to the inhibitor, but the development of more aggressive behavior in a specific environment. As such, this model could provide additional evidence of the disadvantageous use of this compound for the treatment of GBM. It has been reported that IMA enhances GBM invasion through FAK signaling (57). However, we detected an IMA-specific induced activation of GSK3α and GSK3β. GSK3β has been reported to promote invasion in GBM and therapy resistance in cancer (58-60). Therefore, with our results we indicate a potential new resistance mechanism of IMA-induced invasion in GBM.

## CONCLUSIONS

Our study provides relevant information regarding both the MOA and mechanism of resistance following TMZ and sPKI treatments in GBM. As expected from a heterogeneous type of tumor, we showed a diverse biological outcome after prolonged treatment with the same compounds. We demonstrated that this inter-tumoral heterogeneity is ultimately driven by different intracellular network activities orchestrated by kinases, therefore opening horizons to a new strategy to investigate the biological effects of compounds before initiating the clinical phase. Moreover, the detection of the diverse and specific spectrum of kinase activity highlights once more the relevance of pursuing a personalized approach. The application of QuantaKinome™ on the 3D cell cultures models also revealed an ulterior different biological aspect between 2D and 3D drug response in GBM. Therefore, the selection of a suitable model is vital in drug development. As kinase activity is directly linked to the regulation of fundamental biological processes and subsequently to the observed phenotype, future studies are encouraged to pursue the assessment of drug efficacy by coupling the phenotypic evidence with the measurement of kinases activity.

## Author Contributions

SL conceived and designed the study; FF performed the experiments, analyzed and interpreted the data, and wrote the manuscript; NK and EG contributed to the design of the kinomic experiments; NK analyzed the kinomic data; KQ supported to the design and interpretation of the 3D data; NK, EG, KQ, ML, AR, SL supervised the study and revised the manuscript.

## Acknowledgments

We gratefully acknowledge the patients and neurosurgeons who donated and provided the tumor tissues. We express our great appreciation to Ksenia Troshchenkova for her helpful assistance with the QuantaKinome™ analysis. A special thanks goes to Mimetas BV and Pepscope BV for the valuable period spent in their company, and for all the support received.

## Data Availability Statement

Restrictions apply to the availability of the kinomic data, which were used under license for this study. These data supporting the findings of this study are available upon reasonable request to NK at, with the permission of Pepscope BV.

