## Supplementary Figure for "Novel kinome profiling technology reveals drug treatment is patient and 2D/3D model dependent in GBM"

**Supplementary Figures**

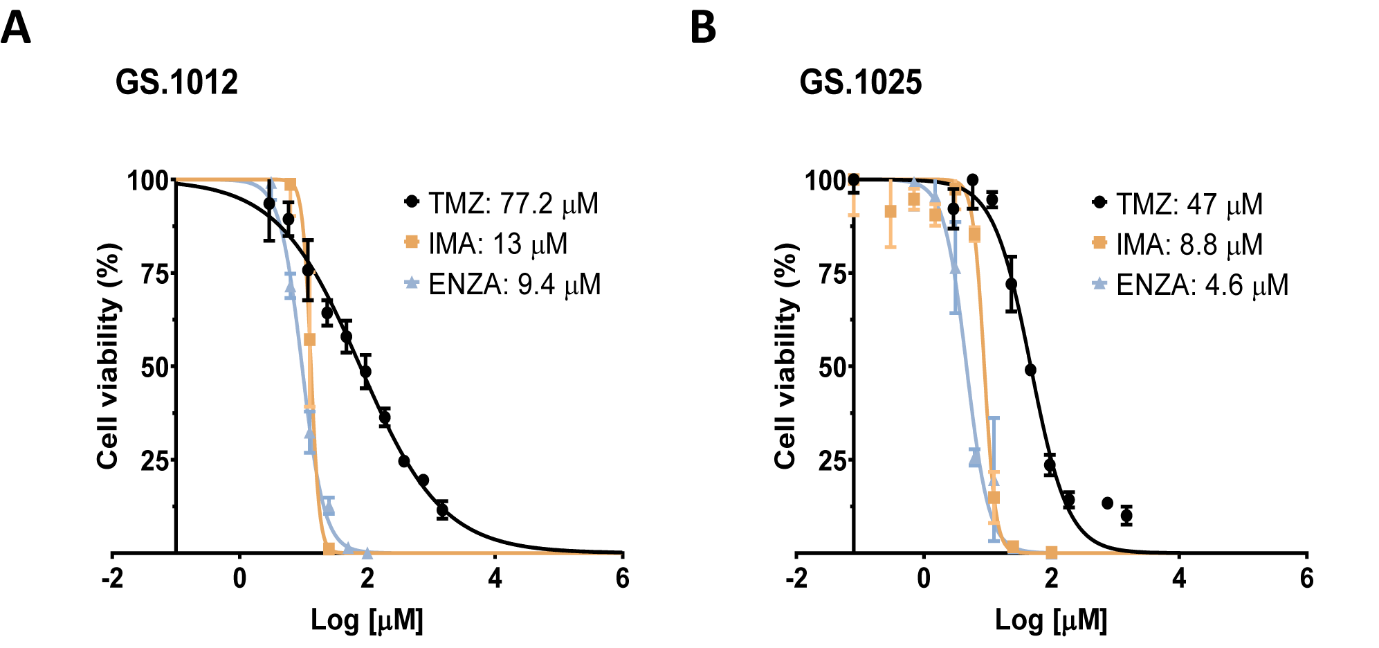

**Supplementary Figure 1.** **Dose-response curves of GS.1012 and GS.1025 treated with temozolomide (TMZ), enzastaurin (ENZA), and imatinib (IMA).
(A)** The dose-response curves in GS.1012 showing the percentage of cell viability versus Log concentration, after 5 days of drug exposure.
**(B)** The dose-response curves in GS.1025 showing the percentage of cell viability versus Log concentration, after 5 days of drug exposure.
Data is presented as mean ± SD (n=3).

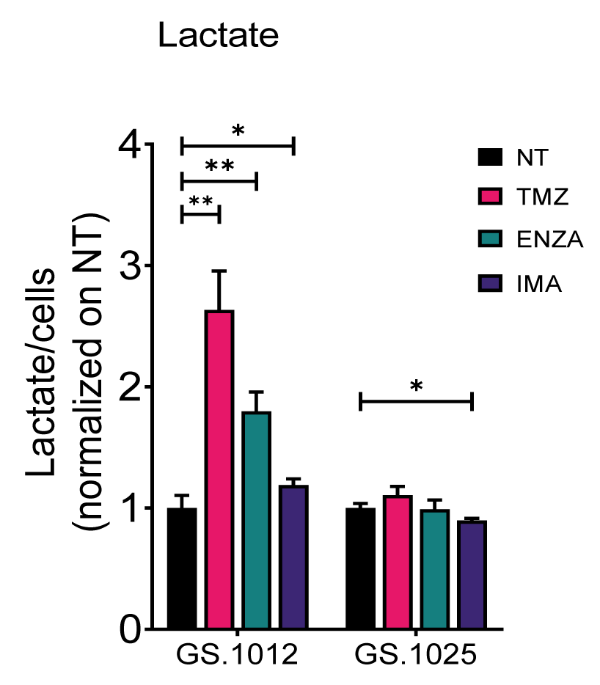

**Supplementary Figure 2. Lactate concentrations of GS.1012 and GS.1025.**
Bar plots showing the lactate concentration in GS.1012 and GS.1025 for temozolomide (TMZ), enzastaurin (ENZA) and imatinib (IMA) treated samples normalized to values observed in untreated (NT) condition. Data are presented as mean ± SD. *p <0.05; **p <0.01

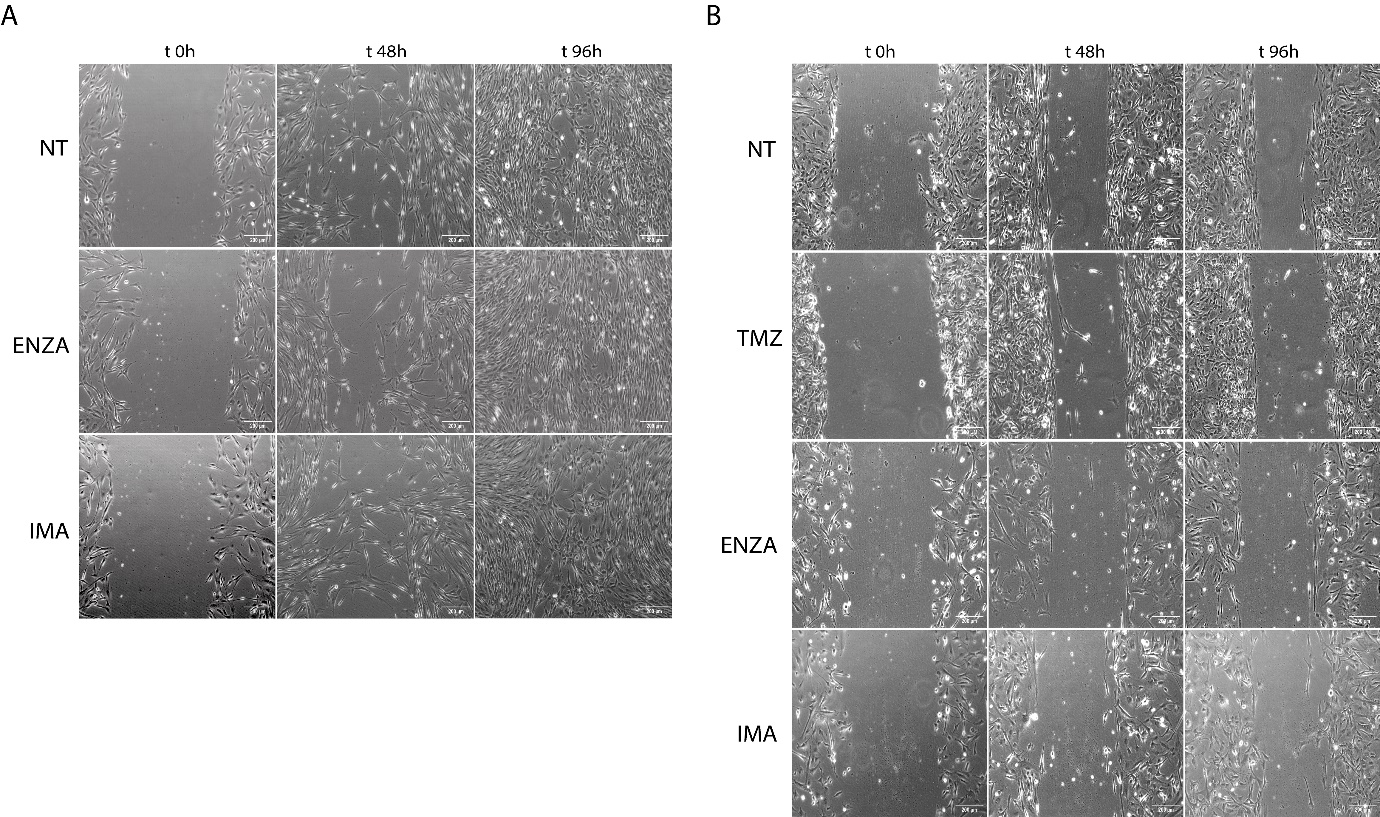

**Supplementary Figure 3**. **Cell migration (scratch migration assay).**
**(A)** Migration of GS.1012 cells was monitored by imaging performed over 96h hours using phase contrast at 10x magnification. Representative images show the untreated control (NT) and cells treated with enzastaurin (ENZA), and imatinib (IMA).
**(B)** Migration of GS.1025 cells was monitored by imaging performed over 96h hours using phase contrast at 10x magnification. Representative images show the untreated control (NT) and cells treated with temozolomide (TMZ), enzastaurin (ENZA), and imatinib (IMA).

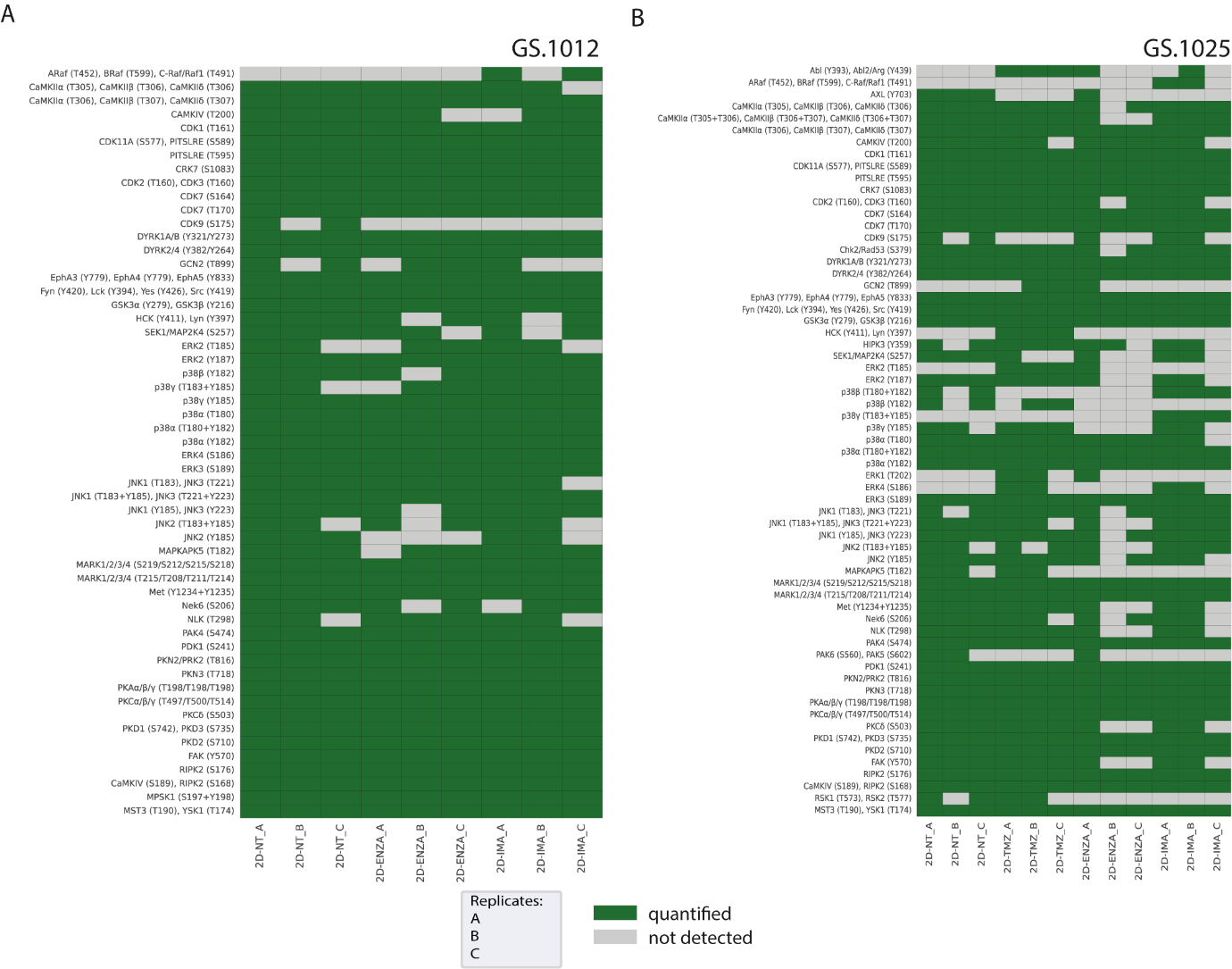

**Supplementary Figure 4**. **Quantified kinase activities in 2D cultures of GS.1012 and GS.1025.
(A)** Map representing in green the quantified kinase activities across 2D cell culture conditions and replicates in GS.1012.
**(B)** Map representing in green the quantified kinase activities across 2D cell culture conditions and replicates in GS.1025.

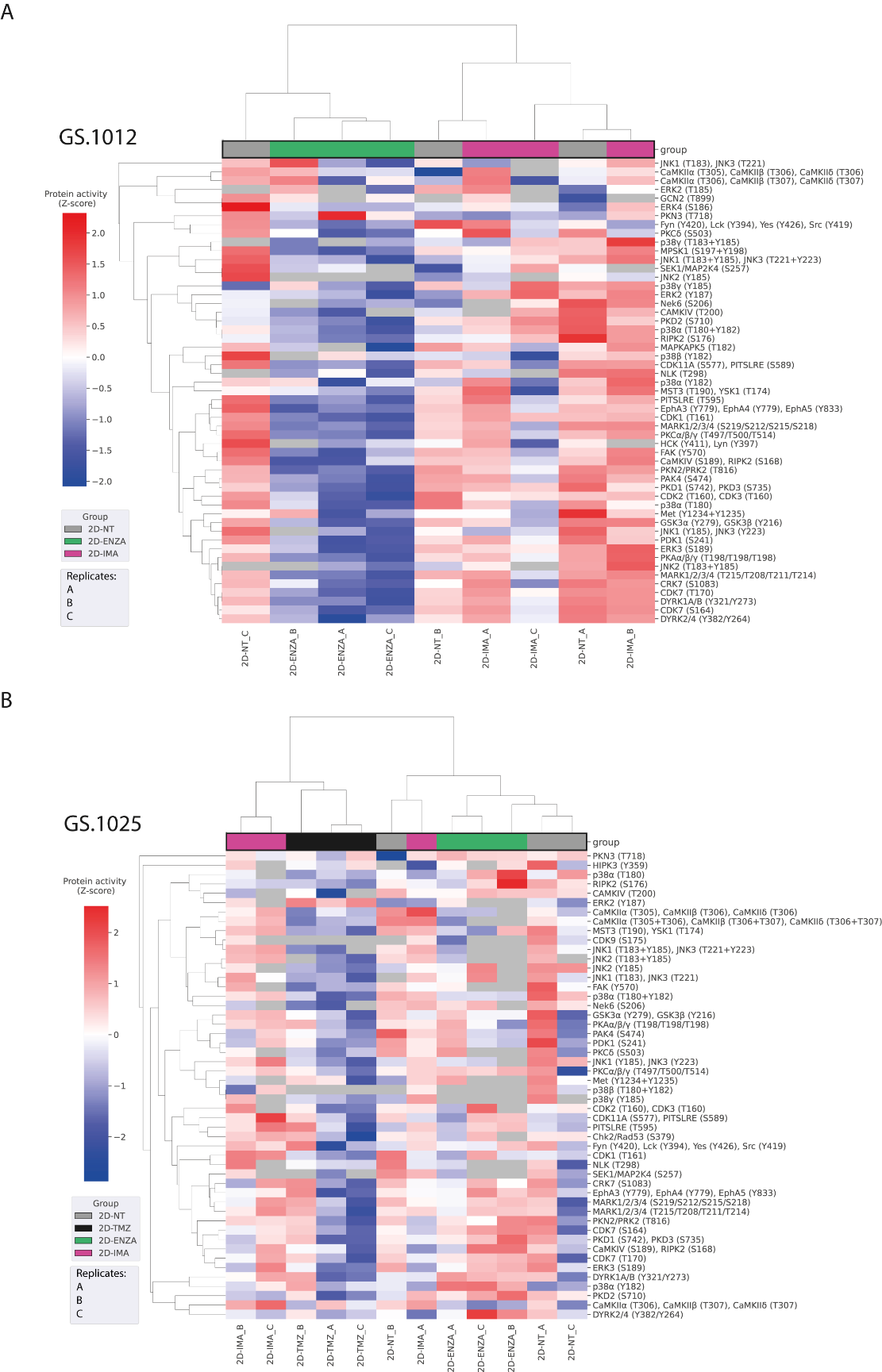

**Supplementary Figure 5**. **Hierarchical clustering of kinase activities in 2D cell cultures.
(A)** Heatmap showing the hierarchical clustering analysis of GS.1012 triplicates treated with temozolomide (TMZ), enzastaurin (ENZA), imatinib (IMA), and the untreated condition (NT).
**(B)** Heatmap showing the hierarchical clustering analysis of GS.1025 triplicates treated with temozolomide (TMZ), enzastaurin (ENZA), imatinib (IMA), and the untreated condition (NT).
Values are expressed as Z-score of the protein activity.

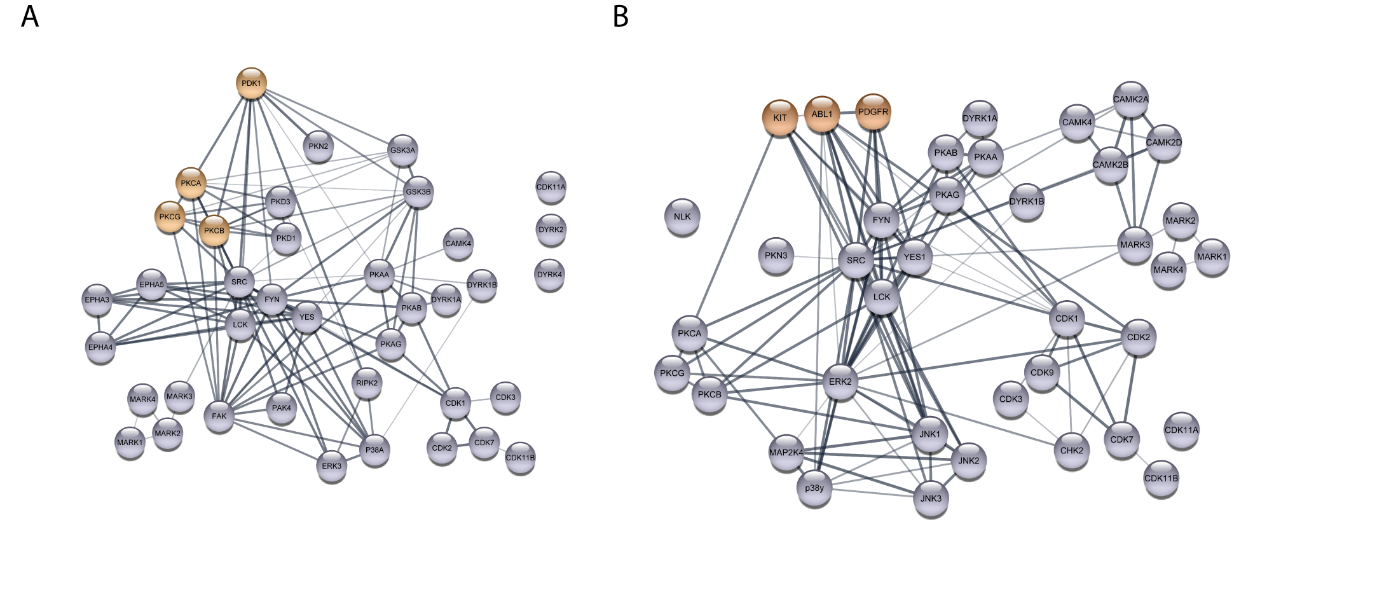

**Supplementary Figure 6. Network analysis of kinase interaction in 2D cell cultures.
(A)** The network represents the interaction among the kinases found significantly affected by enzastaurin treatment in GS.1012. The orange circles represent ENZA targets. The width of the lines represents the interaction score confidence.
**(B)** The network represents the interaction among the kinases with sustained activation after imatinib treatment in GS.1025. The orange circles represent IMA targets. The width of the lines represents the interaction score confidence.

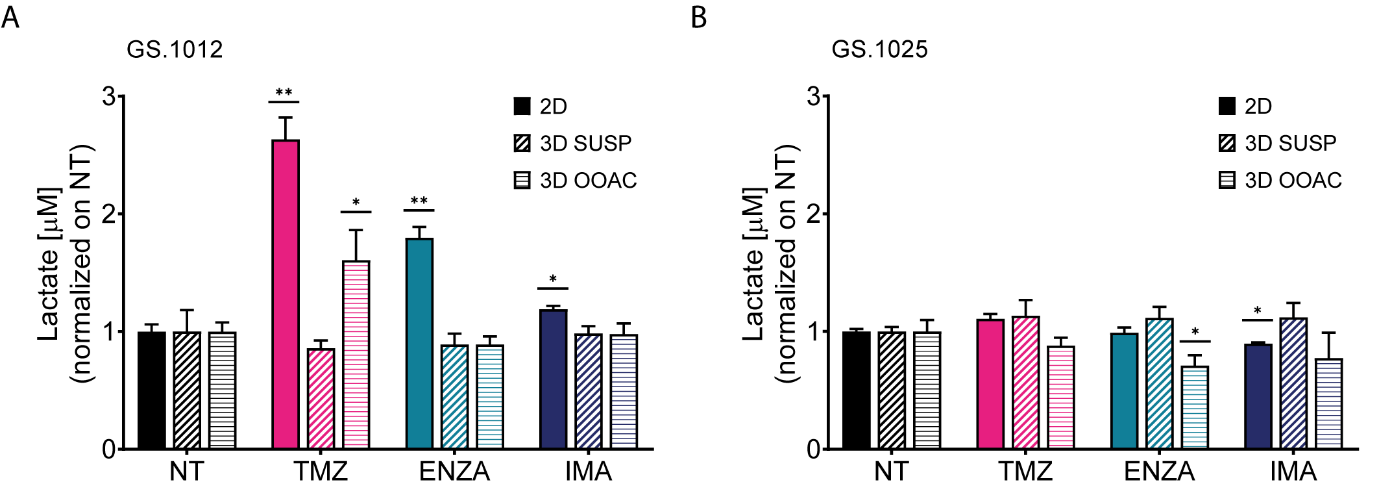

**Supplementary Figure 7. Lactate concentrations of GS.1012 and GS.1025 in 2D and 3D cultures.**
(A) Bar plots showing the lactate concentrations in 2D, 3D-SUSP and 3D-OOAC samples of GS.1012 in temozolomide (TMZ), enzastaurin (ENZA) and imatinib (IMA) treated samples normalized to values observed in non-treated (NT) control.
(B) Bar plots showing the lactate concentrations in 2D, 3D-SUSP and 3D-OOAC samples of GS.1025 in temozolomide (TMZ), enzastaurin (ENZA) and imatinib (IMA) treated samples normalized to values observed in non-treated (NT) control.
Data are presented as mean ± SD. *p < 0.05; **p < 0.01.

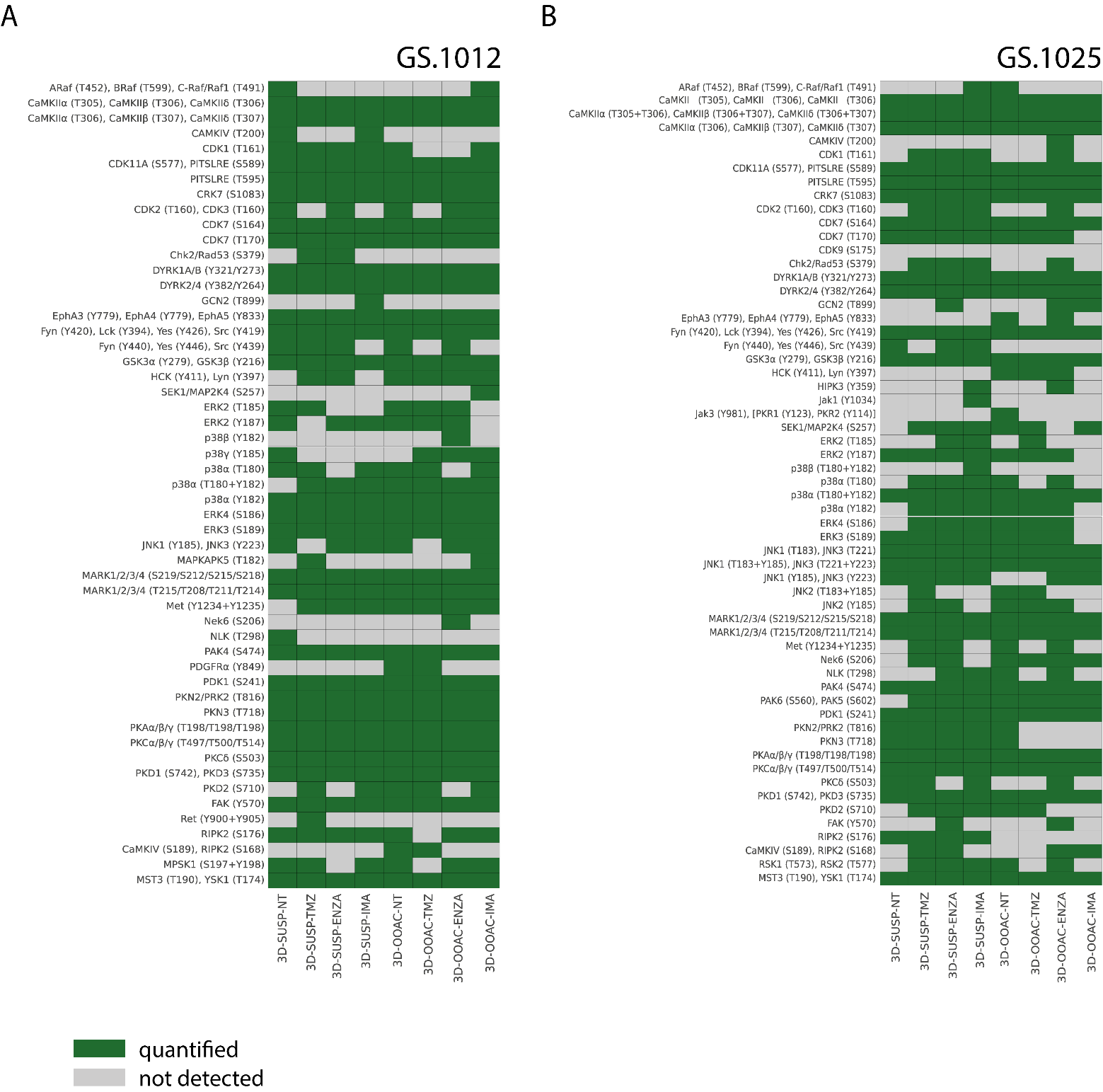

**Supplementary Figure 8**. **Quantified kinase activities in 3D cultures of GS.1012 and GS.1025.
(A)** Map representing in green the quantified kinase activities across 3D-SUSP and 3D-OOAC cell culture conditions in GS.1012. Each sample derives from the pooling of n=22 OMS before quantification.
**(B)** Map representing in green the quantified kinase activities across 3D-SUSP and 3D-OOAC cell culture conditions in GS.1025. Each sample derives from the pooling of n=15 OMS before quantification.

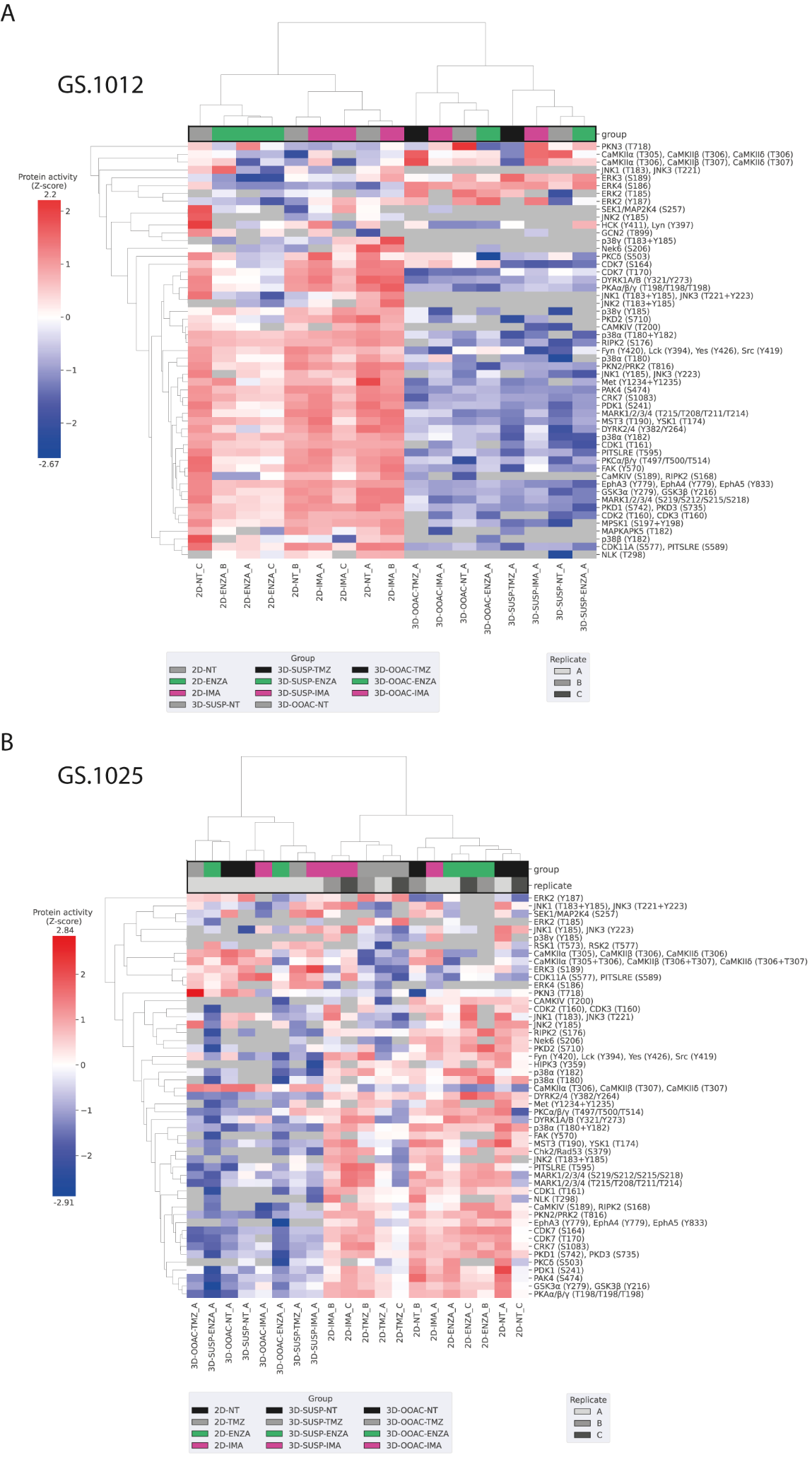

**Supplementary Figure 9. Hierarchical clustering of kinase activities in 2D and 3D OMS.
(A)** Heatmap shows the hierarchical clustering analysis in 2D, 3D-SUSP, and 3D-OOAC samples of GS.1012.
**(B)** Heatmap shows the hierarchical clustering analysis in 2D, 3D-SUSP, and 3D-OOAC samples of GS.1025.
Values are expressed as Z-score of the protein activity.
