## Supplementary Table for "Novel kinome profiling technology reveals drug treatment is patient and 2D/3D model dependent in GBM"

**Supplementary Table 1. Clinical and Molecular Characteristics of Patients**

| Sample | Gender | Age | Diagnosis | Overall survival (months) | Genetic GBM alterations |
| --- | --- | --- | --- | --- | --- |
| GS.1012 | F | 52 | GBM IDH-WT | 3.0 | Amplifications: PDGFRA Homozygous deletion: CDKN2A, CDKN2B Imbalance: Chromosome 7  Loss: Chromosome 10q  Mutations: TERT promoter C228T |
| GS.1025 | M | 71 | GBM IDH-WT | 11.8 | Amplifications: PDGFRA, CDK4 Extra copies: MDM2 Homozygous deletion: CDKN2A Imbalance: Chromosome 7 Loss: Chromosome 10q Mutations: TERT promoter C228T |

**Supplementary Table 2. Size of GS.1012 and GS.1025 OMS in 3D-SUSP and 3D-OOAC.**

| Sample | Condition | 3D-SUSP (mm^2^)  Mean ±SD | 3D-OOAC (mm^2^)  Mean ±SD |
| --- | --- | --- | --- |
| GS.1012 (n=24) | NT | 0.59 ± 0.23 | 0.57 ± 0.21 |
|  | TMZ | 0.52 ± 0.16 | 0.48 ± 0.21 |
|  | ENZA | 0.50 ± 0.17 | 0.40 ± 0.14 |
|  | IMA | 0.42 ± 0.12 | 0.37 ± 0.14 |
| GS.1025 (n=16) | NT | 0.75 ± 0.14 | 0.59 ± 0.18 |
|  | TMZ | 0.74 ± 0.17 | 0.60 ± 0.10 |
|  | ENZA | 0.71 ± 0.20 | 0.60 ± 0.22 |
|  | IMA | 0.77 ± 0.17 | 0.55 ± 0.19 |
